# Endogenous recovery of hippocampal function following global cerebral ischemia in juvenile female mice is influenced by neuroinflammation and circulating sex hormones

**DOI:** 10.1101/2025.01.28.635301

**Authors:** Jose J. Vigil, Erika Tiemeier, James E. Orfila, Nicholas E. Chalmers, Victoria N. Chang, Danae Mitchell, Isobella Veitch, Macy Falk, Robert M. Dietz, Paco S. Herson, Nidia Quillinan

## Abstract

Cardiac arrest-induced global cerebral ischemia (GCI) in childhood often results in learning and memory deficits. We previously demonstrated in a murine cardiac arrest and cardiopulmonary resuscitation (CA/CPR) mouse model that a cellular mechanism of learning and memory, long-term potentiation (LTP), is acutely impaired in the hippocampus of juvenile males, correlating with deficits in memory tasks. However, little is known regarding plasticity impairments in juvenile females. We performed CA/CPR in juvenile (P21-25) female mice and used slice electrophysiology and hippocampal dependent behavior to assess hippocampal function. LTP was and contextual fear were impaired 7-days after GCI and endogenously recovered by 30-days. LTP remained impaired at 30 days in ovariectomized females, suggesting the surge in gonadal sex hormones during puberty mediates endogenous recovery. Unlike juvenile males, recovery of LTP in juvenile females was not associated with BDNF expression. NanoString transcriptional analysis revealed a potential role of neuroinflammatory processes, and specifically Cd68 pathways, in LTP impairment and hormone-dependent recovery. We were able to restore LTP in ovariectomized females with chronic and acute PPT administration, implicating estrogen receptor alpha in recovery mechanisms. This study supports a mechanism of endogenous LTP recovery after GCI in juvenile female mice which differs mechanistically from juvenile males and does not occur in adults of either sex.

## Introduction

Cardiac arrest and consequent global cerebral ischemia (GCI) is a leading cause of death and disability in adults but can also have devastating outcomes in children. Cognitive impairments in survivors of cardiac arrest are observed across all age groups ^1, 2^. Due to advances in resuscitative measures, the survival rate of cardiac arrest (CA) patients has more than doubled in the past decade ^3, 4^, resulting in an increasing population of CA survivors at risk for cognitive dysfunction. Cognitive dysfunction after CA is mainly attributed to widespread hippocampal neurodegeneration induced by the prolonged lack of blood and oxygen to the brain and subsequent excitotoxicity, oxidative stress, and calcium dysregulation ^5^. In addition to the profound neurodegeneration that occurs in the CA1 region of the hippocampus after CA, there are long-lasting deficits in hippocampal-dependent learning tasks and long-term potentiation (LTP), the cellular mechanism of learning and memory^6–8^. Despite extensive effort, neuroprotective strategies aimed at reducing or preventing neuron loss following ischemic insult have not been successfully translated to therapies in humans ^9, 10^. The only available neuroprotective therapy is hypothermia or targeted temperature management, applied in adult and neonatal populations, but this has not been shown to be beneficial in children ^11–13^, highlighting the differences in response to ischemic insult across developmental stages. Clinical evidence also indicates that children and neonates have better odds of survival and more favorable neurological outcome after CA induced global cerebral ischemia compared to adults ^14, 15^. However, there are few studies using preclinical GCI models to evaluate brain function in juvenile populations.

Given the lack of success in translating acute neuroprotective therapies to reduce neuronal cell death ^16^, an alternative strategy is to develop strategies aimed at restoring brain function in surviving neuronal networks, termed *neurorestoration*. Our goal is to identify therapeutic targets to restore the function of the remaining neurons and circuits in the brain to yield a better quality of life for CA survivors. This approach has yielded pre-clinical success in adult animal models, but is understudied in juvenile animals, particularly in females ^17, 18^. Using our translational mouse model of CA and subsequent cardiopulmonary resuscitation (CA/CPR), we have previously shown that there is endogenous recovery of hippocampal function in juvenile male mice after GCI ^19^. Yet, it is not known how long-term hippocampal function in juvenile females is affected by GCI and whether there is similar endogenous recovery of LTP. Therefore, this study utilized our juvenile CA/CPR mouse model in female mice to assess hippocampal function 7- and 30-days after CA/CPR-induced GCI.

Throughout early neurodevelopment, there are profound changes that occur in the mammalian brain such as synaptic pruning, the shift in GABAergic neurotransmission from excitatory to inhibitory ^20^, and the change in NMDAR subunit composition from Glun2B dominant to Glun2A dominant expression ^21^. There are also sex-specific organizational changes that occur in humans during the onset of puberty. The surge of sex hormones, estrogen and progesterone in females and testosterone in males, that occurs during adolescence leads to the development of sex-specific features. Sex hormones, and in particular estrogen, can also have profound effects on synaptic structure, function and plasticity ^22, 23^, but we know little of how the surge of sex hormones during puberty influences the neurological outcome of early-life brain injuries like GCI. Given our previously published results, which revealed endogenous recovery of hippocampal function in juvenile male mice that corresponds with the onset of puberty ^19^, this study evaluates the long-term effect of CA/CPR-induced GCI in juvenile female mice as they mature from juveniles into adults.

Estrogen, one of the major gonadal sex hormones, has been shown to be neuroprotective in many models of neuronal injury ^24, 25^ and there is clinical evidence of better neurological outcome after ischemic insult in pre-menopausal females when compared to similarly aged males or post-menopausal females ^26–28^. Of the female sex hormones produced gonadally, estrogen (E2) is the most prominent and well-studied. E2 elicits its actions in the brain through three known estrogen receptors (ERs) ERα, ERβ, and GPR 30/GPER1 [47, 48]. ERα and ERβ are both expressed in the hippocampus, and both have been shown to impact synaptic transmission and plasticity in adult animals. However, it has been suggested that ERα and ERβ differentially affect learning, memory, and cognition ^29^. While juvenile females have low levels of circulating E2 compared to adult females, a single dose of 17β-estradiol was able to provide neuroprotection after CA/CPR without providing long-term functional benefit ^30^. However, this study did not consider the rapid and sustained increase in E2 that occurs at the onset of puberty, highlighting the need to further study the long-term effect of E2 signaling on GCI outcomes. Therefore, we explored the neurorestorative properties and estrogen receptor specificity of sex hormone signaling to further understand how the surge of gonadal sex hormones at the onset of sexual maturity in juvenile female mice influences hippocampal function after CA/CPR-induced global ischemia.

GCI was induced in pre-pubescent female mice and measurements of hippocampal function (LTP and contextual fear conditioning) and transcriptional alterations were made at acute (pre-puberty) and chronic (post-pubescent) time points after GCI. We then altered endogenous sex hormone signaling through ovariectomy and hormone replacement to assess the neurorestorative capacity of gonadal hormone signaling on hippocampal function. We identify a critical role for circulating sex hormones and implicate potential molecular pathways that underlie plasticity deficits and estrogen-dependent endogenous recovery of LTP.

## Materials and Methods

### Animals

All experimental protocols were approved by the Institutional Animal Care and Use Committee (IACUC) at the University of Colorado, Anschutz Medical Campus and adhered to the National Institute of Health guidelines for the care and use of animals in research. Female C57/BL6 mice were PND 21-25 at time of cardiac arrest and subsequent resuscitation. All mice were permitted free access to water and standard lab chow ad libitum with a 14/10-h light/dark cycles. If mice were littermates, they were equally divided between experimental groups.

### Cardiac Arrest/Cardiopulmonary Resuscitation

As we have previously described for the juvenile CA/CPR mouse model ^31^, female P21-25 (developmentally equivalent to 1-5 year old child ^32^) Charles River C57BL/6 mice were anesthetized using 3% isoflurane and intubated with 20% oxygen (100cc/min) 80% air mixture (900cc/min). Respiration was set to 160 strokes/minute with a stroke volume of 80-150µL (calculated as eight times mouse’s body weight in grams with a maximum of 150 µL). A jugular catheter was inserted, and electrocardiogram (ECG; Corometrics Eagle 4000N., Milwaukee WI) electrodes connected to display heart rate. Using a heat lamp and water-filled head coil, body and head and temperatures were brought to 37.5±0.5°C. To initiate cardiac arrest (CA), 0.05mL 0.5mEq/mL potassium chloride was injected via jugular catheter, respirator was disconnected, and asystole confirmed for 8 minutes via ECG readout. During CA, head temperature was maintained at 37.5±0.5°C using a 40°C water-heated head coil while body temperature was permitted to fall to a minimum temperature of 35.5°C. Mechanical ventilation (MiniVent Type 845., Germany) was initiated 30 seconds prior to resuscitation at a rate of 210 breaths/minute with 100% oxygen at 300cc per minute. Resuscitation was initiated via chest compressions and up to 0.5mL 16µg/mL epinephrine administered via the jugular catheter over a period of 3 minutes until normal sinus rhythm was achieved. After reaching a recovery of spontaneous respiration rate of 30 breaths per minute, air was increased to 300cc/minute and respiration rate reduced by 10 strokes per minute for 5 minutes until extubating. Animals were excluded if return of spontaneous circulation was not achieved within 3-minutes of initiation compressions or if return of spontaneous breathing with successful extubation was not achieved within 30 minutes after resuscitation. Mice were single housed after recovery from anesthesia and post-surgical care consisted of 2-3 days of housing cages on a warming pad, provision of soft chow, and daily administration of 0.5mL subcutaneous 0.9% saline.

### Open Field Testing

#### Apparatus

The open field (OF) apparatus consisted of a square white chamber (14” L x 14” W x 13”H) with an open top. A camera was mounted above each chamber to record the activity of the mice during the test. The chamber was divided into two zones, an inner zone was indicated with permanent marker on the chamber floor for consistent placement of measurement zones within the recording and analysis software, Any-maze (Stoelting, Wood Dale, IL). The outer zone was the rest of the chamber surrounding the inner zone. A small lamp was placed in the behavior room to ensure a consistent lighting level (∼20 lux) and to ensure there were no shadows cast into the chamber. The chamber was cleaned with 70% isopropanol between animals.

#### Procedure

Seven- or thirty-days post-CA/CPR, juvenile female (PND 28-32 or PND 51-54) mice were subjected to the OF test to assess anxiety like behavior and locomotor activity. Animals were habituated to the behavior room in their home cage for 1-hour before beginning the testing procedure. At the start of testing, an animal was placed into the chamber and video recording was begun in Any-Maze (Stoelting, Wood Dale, IL). The animal was allowed to explore the apparatus for 10 minutes before being returned to its home cage. Time spent in the inner and outer zone was assessed by the software and percent time spent in the open (center zone) was calculated by dividing the amount of time spent in the center of the apparatus by the total time of the test and multiplying by 100. Average speed and total distance traveled was also calculated within Any-maze (Stoelting, Wood Dale, IL) for each animal.

### Contextual Fear Conditioning

#### Apparatus

The fear conditioning apparatus consisted of a sound attenuating chamber (24” L x 14” W x 14”H) which contained a grid-shock floor (Coulbourn, Holliston, MA) onto which a 5-sided plexiglass cube (6 x 6 x6) was placed to contain the mice during the experiments (Context A). The shock floor was connected to a shock stimulator (Coulbourn, Holliston, MA) that was automatically triggered by an Arduino control board. A foot-shock (0.5mA, 1s) delivered through the floor was used for the unconditioned stimulus (US). For all procedures carried out in Context A, the apparatus was cleaned with a 70% isopropanol solution before and after each animal, and each animal was transported to the conditioning room in their home-cage. Any-maze software (Stoelting, Wood Dale, IL) was used to record and assess freezing behavior during every stage of the procedure.

#### Procedure

7- or 30- days post-CA/CPR, juvenile (PND 28-32 or PND 51-54) female mice were subjected hippocampal dependent contextual fear conditioning. On the first day of the procedure, animals were habituated to the conditioning room while in their home cage for one-hour. Animals were also habituated to Context A for 5-minutes without presentation of the US. 24-hours later, animals were re-exposed to Context A to obtain a 2-minute baseline before delivering foot shocks. After the baseline period, three US were delivered with 1-minute separating each pairing. After delivery of the last US, mice were allowed to remain in the chamber for 1 minute and 45 seconds before being returned to their home cage. The next day, animals were returned to context A for 5-minutes without presentation of the US to assess freezing behavior as a measure to learn and remember contextual information.

### Ovariectomy and Osmotic Pump Implantation

As previously described ^33^, ovariectomy (OVX) was performed 5-7 days post CA. Mice were anesthetized using 3% isoflurane and intubated with 20% oxygen (100cc/min) 80% air mixture (900cc/min) and placed in sternal recumbency. A 0.05 mL bolus 0.25% bupivacaine was administered subcutaneously and 15mm incision made parallel to spine near lumbar portion of the back. The skin was separated from the muscle, and a 2mm incision made through the external obliques where they meet the dorso-lumbar subcutaneous fat and gluteus superficialis. Forceps were used to extract the ovary and fat pad and a hemostat was placed below the junction of oviduct and uterus. Ovaries were severed and hemostats were released. In animals receiving hormone replacement, after removal of both ovaries, a subcutaneous pellet or osmotic pump (Alzet; Cupertino, CA) was inserted into the lower lumbar space, and the incision was closed with surgical staples. Osmotic pumps contained either Vehicle (DMSO in saline), E2 (17β-estradiol equivalent), DPN (ERβ agonist), or PPT (ERα agonist), and delivered each drug at a rate of 1mg/kg/day. For acute PPT administration in gonadally intact females, IP injections were performed at 6 days after CA/CPR.

### Hippocampal Slice Preparation

Hippocampal slices were prepared at 7- or 30-days after recovery from CA/CPR or sham surgeries as previously reported ^31^. Mice were anesthetized with 5% isoflurane in an O2-enriched chamber and transcardially perfused with ice-cold (2–5 °C) oxygenated (95% O2/5% CO2) artificial cerebral spinal fluid (aCSF) for 2 min prior to decapitation. The brains were then rapidly extracted and sectioned in ice-cold aCSF. The composition of aCSF was the following (in mM): 126 NaCl, 2.5 KCl, 25 NaHCO3, 1.3 NaH2PO4, 2 CaCl2, 1 MgCl2, and 10 glucose. Transverse hippocampal slices (300 μm thick) were cut with a Vibratome VT1200S (Leica., Deer Park IL) and transferred to a holding chamber containing room temperature aCSF for at least 1 h before recording.

### Extracellular Field Potential Recordings

Synaptically evoked field potentials were recorded from slices containing the hippocampal CA1 region, which were placed on a temperature controlled (31 ± 0.5 °C) interface chamber perfused with aCSF at a rate of 1.5 mL/min. Field excitatory post-synaptic potentials (fEPSP) were produced by stimulating the Schaffer collaterals (SC = CA3 axons) and recording in the stratum radiatum (SR) of the CA1 region. Analog fEPSPs were amplified (1000×) and filtered through a preamplifier (Model LP511 AC, Grass Instruments., West Warwick RI) at 1.0 kHz, digitized at 10 kHz and stored on a computer for later off-line analysis (Clampfit 10.4, Axon Instruments). The rise slope of the fEPSP was measured and adjusted to 50% of the maximum slope and test pulses were evoked every 20 s. Paired-pulse responses were recorded using a 50-ms inter-pulse interval (20 Hz) and expressed as a ratio of the slope of the second pulse over the first pulse. A 20-min stable baseline was established before delivering a theta burst stimulation (TBS) train of four pulses delivered at 100 Hz in 30-ms bursts repeated 10 times with 200-ms inter-burst intervals. Following TBS, the fEPSP was recorded for an additional 60 min. The averaged 10-min slope from 50 to 60 min after TBS was divided by the average of the last 10-min of the baseline (set to 100%) prior to TBS to determine the amount of potentiation. For time course graphs, normalized fEPSP slope values were averaged and plotted as the percent of baseline. A maximum of two slices per animal were used for analysis (n = number of slices).

### NanoString

RNA was isolated from flash frozen whole mouse hippocampi using an RNA-Aqueous PCR kit (Invitrogen., Waltham MA). Briefly, tissue was homogenized in lysis/binding solution in an Eppendorf tube with a plastic pestle. An equal volume of ethanol (64%) was added to the tubes and vortexed. Contents of each tube were then transferred to a filter cartridge inside of a collection tube and centrifuged at 10,000-15,000 x g for 30s. The tubes were washed with wash solution #1 and centrifuged at 10,000-15,000 x g for 30s and washed with solution #2/3 twice with centrifugation at 10,000-15-000 x g. mRNA was then eluted from the tube with preheated elution solution and stored at −80°C until performing the NanoString procedure. A Bioanalyzer was used to determine RNA concentrations and integrity. Sample with RNA integrity values below 8 were not included in NanoString analysis.

Samples were diluted to 5ng/µl, and NanoString (NanoString Technologies Inc., Seattle WA) runs were performed by the University of Colorado-Anschutz Veteran’s Affairs Research Core using the NanoString N-counter. Every sample was obtained from an individual animal and are thus independent replicates. Each sample was run using the Neuropath panel which contained 10 reference housekeeping genes (Aars, Asb7, Ccdc127, Cnot10, Csnk2a2, Fam104a, Lars, Mto1, Supt7l, Tada2b), 8-negative controls, and 6-positive controls. All data passed quality control checks, with no normalization flags. Data was analyzed by ROSALIND® (https://rosalind.bio/), with a HyperScale architecture developed by ROSALIND, Inc. (San Diego, CA). Normalization, fold changes and p-values were calculated using criteria provided by NanoString. ROSALIND follows the nCounter Advanced Analysis protocol of dividing counts within a lane by the geometric mean of the normalizer probes from the same lane. Fold changes and p values were calculated using the fast method as described in the nCounter® Advanced Analysis 2.0 User Manual^34–38^. P-value adjustment is performed using the Benjamini-Hochberg method of estimating false discovery rates (FDR). A protein-protein interaction plot was prepared using the online analysis software STRING and lollipop plot was generated using R software (Version:2022.07.2; Packages: DOSE, ggplot2, forcats, org.Mm.eg.db, enrichplot).

### Immunohistochemistry

7- or 30-days post-CA/CPR or sham surgery, juvenile (PND 28-32 or PND 51-54) female mice were transcardially perfused with ice cold phosphate buffered saline (PBS) for 5 minutes followed by 5 minutes of ice cold 4% paraformaldehyde (PFA). The brains were then rapidly extracted and placed in PFA overnight at 4°C. The next day, the brains were placed into cryoprotection solution (20% glycerol, 20% Sorenson’s Buffer, 60% water) at 4°C until the brain sank to the bottom of the vial (overnight). The cryoprotected brains were then stored at 4°C until slicing of the brain. Brains were sliced in the coronal orientation (30µm) on a sliding microtome (Leica SM 2010R, Germany) using OTC compound and crushed dry ice to secure the brain to the cutting stage and freeze the brain itself. Slices were collected with a small paint brush and placed into a 24-well plate containing cryostorage (50% 0.2M PO4, 30% ethylene glycol, 1% polyvinylpryyolidine, 19% sucrose) solution until processing for immunohistochemistry. Free-floating sections were washed 3x for 5-minutes each in Phosphate Buffered Saline (PBS) before blocking in PBS-Triton X (PBST)solution containing 5% Normal Donkey Serum (NDS) for 1-hour at room temperature. Slices were then incubated in primary goat anti-mouse Iba1 polyclonal antibody (Abcam, Waltham, MA) and rat anti-mouse CD68 monoclonal antibody (Bio-rad, Hercules, CA) at 1:500 in 3% PBST solution overnight at 4°C. Slices were then washed 3x in PBS for 5-minutes each and incubated in donkey anti-goat (Jackson ImmunoResearch, Westgrove, PA) secondary antibody conjugated to AlexaFluor 594 and goat anti-rat (Jackson ImmunoResearch, Westgrove, PA) secondary conjugated to AlexaFluor 488, both at 1:500 in PBST containing 3% NDS for 1-hour at room temperature. Sections were then washed 3x in PBS for 5-minutes before mounting onto charged slides and being cover slipped in DPX (Sigma, St. Louis, MO) mounting media. Confocal images were obtained using an Olympus FV1000 laser scanning confocal microscope, 40x oil immersion objective with Fluoview software (Olympus FluoView, FV10-ASW). Images of the hippocampal CA1 region were attained with step sized of 0.1µM at a resolution of 1024 × 1024. A single maximum intensity z-projection was acquired in two consecutive hippocampal sections in the coronal orientation and from each hemisphere, yielding a total of four images per animal. The integrated density of fluorescence was then acquired in Image J (NIH) for each channel (red-Iba1; green-CD68) by placing four equally sized regions of interest (ROIs) onto the maximum intensity projection. Two ROIs were place onto the synaptic layer (SC-CA1) and two ROIs were place onto the cellular layer (CA1) for each image. The integrated density values for CD68 fluorescence were normalized to CD68 fluorescence levels in 7-day shams for comparison. Data is visualized by plotting the average normalized integrated density per animal (n = # of animals) and expressed as mean ± SD. However, the expression of CD68 was statistically compared with a general linear mixed model (GLMM), given the multiple measurements taken from each animal.

### Rigor and Statistics

To enhance the rigor and reproducibility of our study, all experimenters were blind to injury status of animals and drug treatment through data acquisition. All experiments followed ARRIVE 2.0 guidelines ^39^. Data was compiled and grouped for statistical comparison after completion of all data acquisition to limit experimenter bias. All animals were randomly assigned to experimental groups by a surgeon who performed all CA/CPR, OVX, and osmotic pump surgeries and a separate experimenter performed all acquisition and analysis of data to ensure blinding. Sample size and power analysis were performed with previous data generated in our laboratory using G-power and group sizes of 8 were needed to detect 20% change with 90% power for LTP experiments. All data are expressed as mean ± SD unless otherwise stated. Normality of data was assessed with the Anderson-Darling normality test^40^ to ensure the proper statistical tests were performed. Where appropriate, either a one-way ANOVA, GLMM, or Student’s T-Test was performed to assess statistical significance. Differences were considered statistically significant if p < 0.05.

## Results

### Hippocampal Long-term Potentiation is Acutely Diminished and Endogenously Recovers after CA/CPR Induced GCI in Juvenile Female Mice

Long-term potentiation (LTP) in the hippocampus has been strongly associated with cognitive function^41–45^. This cellular mechanism of learning and memory can be assessed in acute hippocampal brain slices with *ex vivo* electrophysiological recordings of field excitatory postsynaptic potentials (fEPSP). To evaluate hippocampal plasticity after juvenile GCI, LTP recordings were performed either 7- or 30- days after recovery from CA/CPR or sham surgery. LTP recordings in sham mice 7-days after surgery displayed robust potentiation of the Schaffer Collateral (SC) to CA1 synapse (Figure 1A) after theta burst stimulation (TBS), which remained potentiated until the end of the recordings, 60-minutes later (145.3±21.77% of BL, n=10 (Figure 1B, C). In contrast, 7-days after CA/CPR there was significantly diminished hippocampal LTP (115.4±22.75% of BL; p < 0.05 compared to sham, N=12) with a concomitant reduction of the paired pulse ratio (PPR) (sham: 1.73±0.2, n=7; 7-day CA: 1.47±0.08, n=9, p<0.01) (Figure 1C). The decreased PPR is indicative of an increased presynaptic probability of glutamate release ^46^. Interestingly, there was endogenous recovery and no evidence of diminished LTP by 30-days after CA/CPR (143.2±22.56% of BL, n=10, p>0.05 compared to sham and p<0.05 compared to 7-day CA/CPR). While LTP recovered at 30-days, PPR remained reduced (1.445±0.12, n=12) compared to sham (p<0.001) (Figure 1C). The altered release probability in this study prompted us to determine if LTP in this circuit is purely post-synaptic, so we also performed LTP studies in sham and CA/CPR animals in the presence of AP-5. As expected, LTP cannot be induced in this circuit when AP-5 is included in the ACSF (Supplemental Figure 1). These results suggest that LTP impairments and reduced paired pulse ratio are perhaps unrelated phenomenon with distinct underlying mechanisms.

**Figure 1:**
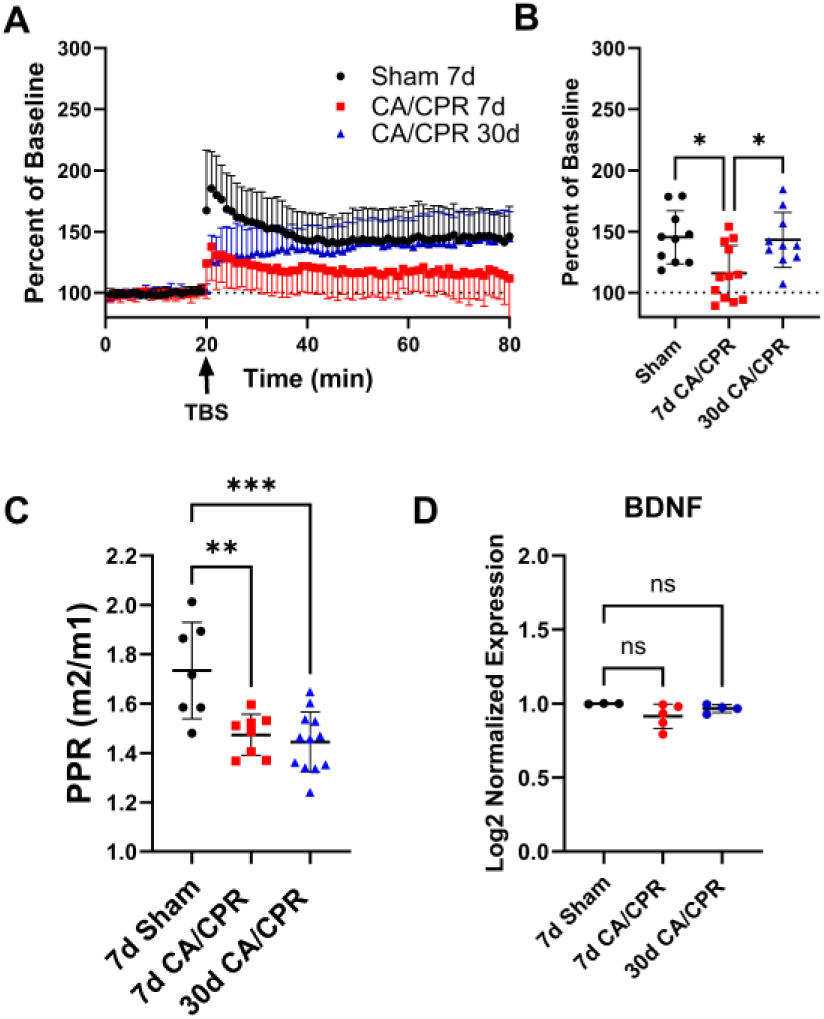
In female mice, CA/CPR induced GCI results in diminished hippocampal LTP that endogenously recovers within 30-days. (A) Hippocampal LTP recordings reveal diminished LTP 7-days after CA/CPR which recovers to sham levels by 30-days after CA/CPR. (B) Quantification of data in (A), using the last 10-minutes of the recording for comparison between groups. (* = p < 0.05, One-way ANOVA). (C) Paired Pulse Ratio measurements reveal a CA/CPR induced increase in synaptic transmission (reduced PPR) that remains despite recovery of Hippocampal LTP (** = p < 0.01, *** = p < 0.001, One-way ANOVA). (D) Hippocampal BDNF mRNA expression is not altered by GCI. (ns = p > 0.05, One-way ANOVA).

### Hippocampal Dependent Learning and Memory is Acutely Diminished and Endogenously Recovers After CA/CPR Induced GCI in Juvenile Female Mice

To corroborate the results observed in Figure 1, we performed contextual fear conditioning experiments in sham and CA/CPR animals at acute (7D) and chronic (30D) time points after surgery. The hippocampus is heavily involved in this sort of spatial learning and memory task^47^. So, to support the role that hippocampal LTP has in learning and memory, we performed contextual fear conditioning and assessed locomotor function, to ensure that locomotor activity is not influencing freezing behavior measured during fear conditioning. We found that 7-days after sham or CA/CPR surgery there is no change in locomotor function as there is no significant difference in the distance traveled (Sham: n = 8, 18.26 ± 3.79 m; CA/CPR: n = 6, 20.45 ± 9.49m; p > 0.05) (Figure 2A) or average speed (Sham: n = 8, 0.061 ± 0.012 m/s; CA/CPR: n = 6, 0.068 ± 0.031 m/s; p > 0.05) (Figure 2B) during testing in the OF. There was also no evidence of anxiety like behavior, as sham animals (n = 8, 59.74 ± 14.98% of time in center) spent similar amounts of time in the center of the apparatus as CA/CPR animals (n = 6, 37.14 ± 20.5 % time in center, p > 0.05) (Figure 2C). There was, however, a significant decrease in the amount of time spent freezing upon re-exposure to context A, after contextual fear conditioning in CA/CPR animals (59.74 ± 14.98% freezing, n = 6), compared to shams (37.14 ± 20.5% freezing, n = 8, p < 0.05) (Figure 2D). 30-days after surgery there was also no difference in distance traveled (Sham: n = 7, 22.24 ± 4.29m; CA/CPR: n = 7, 23.39 ± 6.13m; p > 0.05) (Figure 2E), average speed (Sham: n = 7, 0.074 ± 0.014 m/s; CA/CPR: n = 7, 0.078 ± 0.021 m/s; p > 0.05) (Figure 2F), or the presence of anxiety-like behavior (Sham: n = 7, 16.67 ± 6.58% time in center; CA/CPR: n = 7, 12.16 ± 7.23% time in center) (Figure 2G). However, in agreement with the LTP findings, we found that the acute impairment of hippocampal-dependent contextual fear observed above, endogenously recovers by 30-days after CA/CPR. There is no significant difference between sham (n = 7, 63.61 ± 19.43% freezing) and CA/CPR animals (n = 7, 54.31 ± 25.35% freezing; p > 05) (Figure 2H) in the amount of time spent freezing in an aversive context, 30-days after surgery. Thus, in addition to the acute disruption and endogenous recovery of hippocampal LTP after GCI, there is also a corresponding deficit in hippocampus dependent learning and memory that recovers by 30-days after CA/CPR.

**Figure 2:**
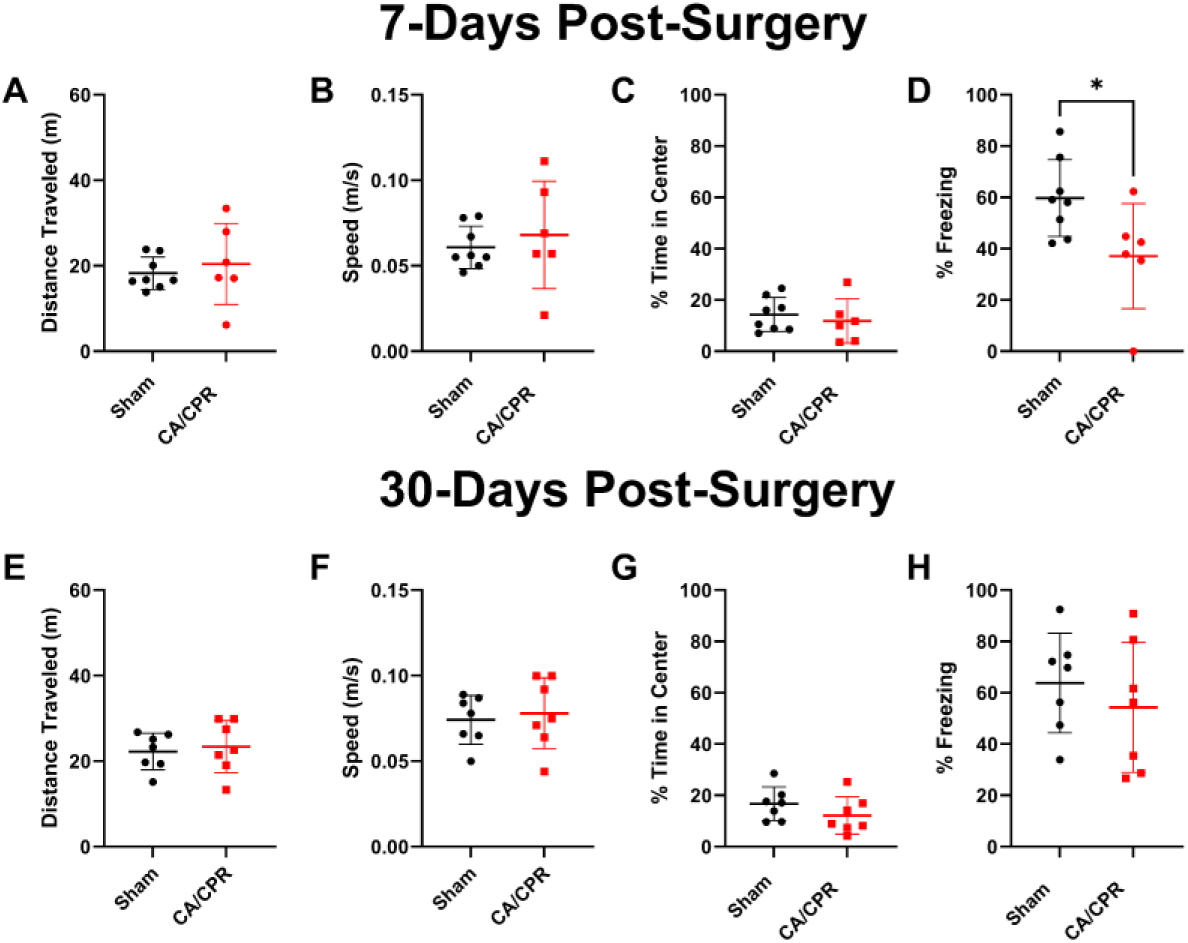
In female mice, CA/CPR induced GCI results in spatial learning and memory deficits that endogenously recover within 30-days. (A-D = 7D after surgery). (A) Distance traveled in the OF test. (B) Average speed in the OF test. (C) % time in center of OF apparatus. (D) Percent of time spent freezing when re-exposed to the aversive context A. (E-H = 30D after surgery). (E) Distance traveled in the OF test. (F) Average speed in the OF test. (G) % time in center of OF apparatus. (H) percent of time spent freezing when re-exposed to the aversive context A. (* = p < 0.05, Students T-test).

### Enhanced Expression of Neuroinflammatory Transcripts Likely Mediates the GCI induced Deficit of Hippocampal LTP

To gain deeper insight into the mechanism of hippocampal LTP deficits and endogenous recovery, we performed transcriptional analysis using NanoString technology neuropathology panel in whole hippocampus. Given our previous findings that LTP impairment and endogenous recovery in males was correlated with BDNF, we first compared BDNF transcript levels in females. Surprisingly, we saw no changes in BDNF expression between groups (Relative to 7d Sham (n = 3), 7d CA/CPR = 0.92±0.002 (n = 5); 30d CA/CPR = 0.97±0.083 (n = 3); p > 0.05) (Figure 1D). Despite the lack of effects on BDNF expression, we were able to identify 151 transcripts that were significantly upregulated and 11 transcripts that were significantly downregulated by CA/CPR after 7-days (n = 5), relative to age-matched sham animals (n = 3) (Figure 3A top left,). In contrast, there were 5 transcripts that were significantly upregulated and 103 transcripts that were significantly downregulated by CA/CPR after 30-days of recovery (n = 4), relative to 7-day CA/CPR animals (n = 3) (Figure 3A top right). Thus, we sought to identify common transcripts that were upregulated at 7-days post CA/CPR and downregulated at 30-days post CA/CPR. These transcripts may underlie the functional hippocampal LTP deficit and correspond with endogenous recovery after GCI. There was a total of 78 transcripts that were upregulated 7-days post CA/CPR and recovered by 30-days post CA/CPR (Figure 3A bottom). Of these 78 transcripts, there were 31 with greater than 2-fold increase at 7-days post CA/CPR compared to sham animals and many of these had a similar 2-fold decrease by 30-days post CA/CPR (Figure 3B) when compared to 7-day CA/CPR. Using those 31 transcripts, a protein-protein interaction analysis was performed, revealing CD68, Spi1, Trem2, C1qa, Apoe and Cx3cr1 as some of the central mediators of the response to GCI induced by CA/CPR (Figure 3C), all of which are involved in neuroinflammation. By 30-days post CA/CPR, these same neuroinflammatory pathways were negatively regulated as revealed by gene ontology analysis (Figure 3D). Thus, our results suggest that there is an acute inflammatory response to GCI in the juvenile female hippocampus associated with LTP and spatial learning impairments. As GCI-induced neuroinflammatory gene expression subsides at 30 days post-CA/CPR, there is a concomitant recovery of hippocampal LTP and hippocampal dependent learning and memory (Figure 1 and Figure 2).

**Figure 3:**
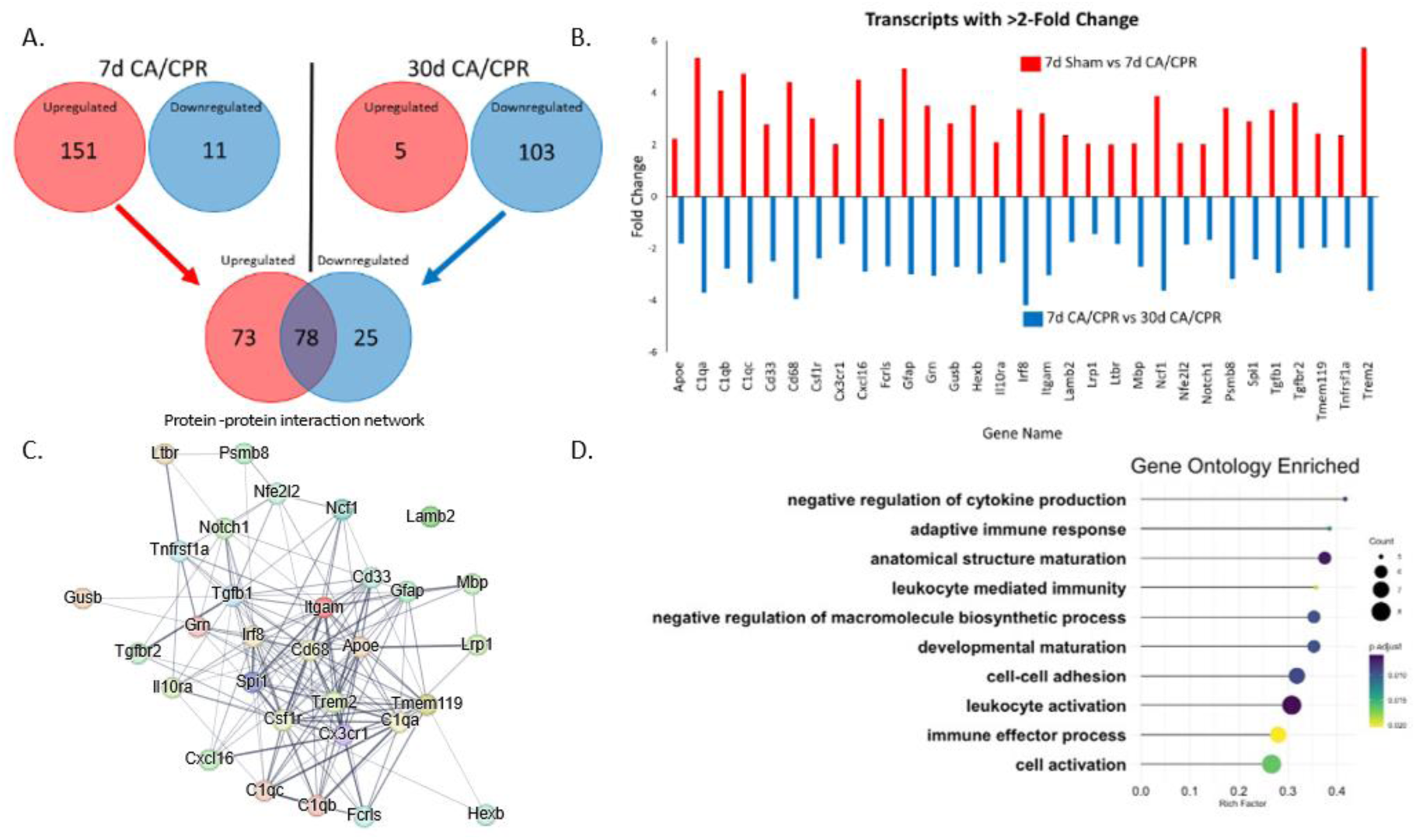
Transcriptional analysis of the observed recovery of hippocampal function in Figure 1. (A) Venn diagram depicting the number of transcripts differentially expressed 7-days after CA/CPR (left) or 30-days after CA/CPR (right). Lower Venn diagram depicts the number of transcripts that were altered by CA/CPR at 7-days and recover by 30-days. (B) Bar graph depicting a total of 31 transcripts from (A) had greater than 2-fold change in expression. (C) STRING protein-protein interaction network of the 31 transcripts in (B) revealing Cd68, Spi1, and Trem2 as a central regulator of the altered transcripts. (D) Lollipop plot depicting the 10 pathways that are most affected by CA/CPR and subsequent recover by 30-days after CA/CPR.

**Figure 4:**
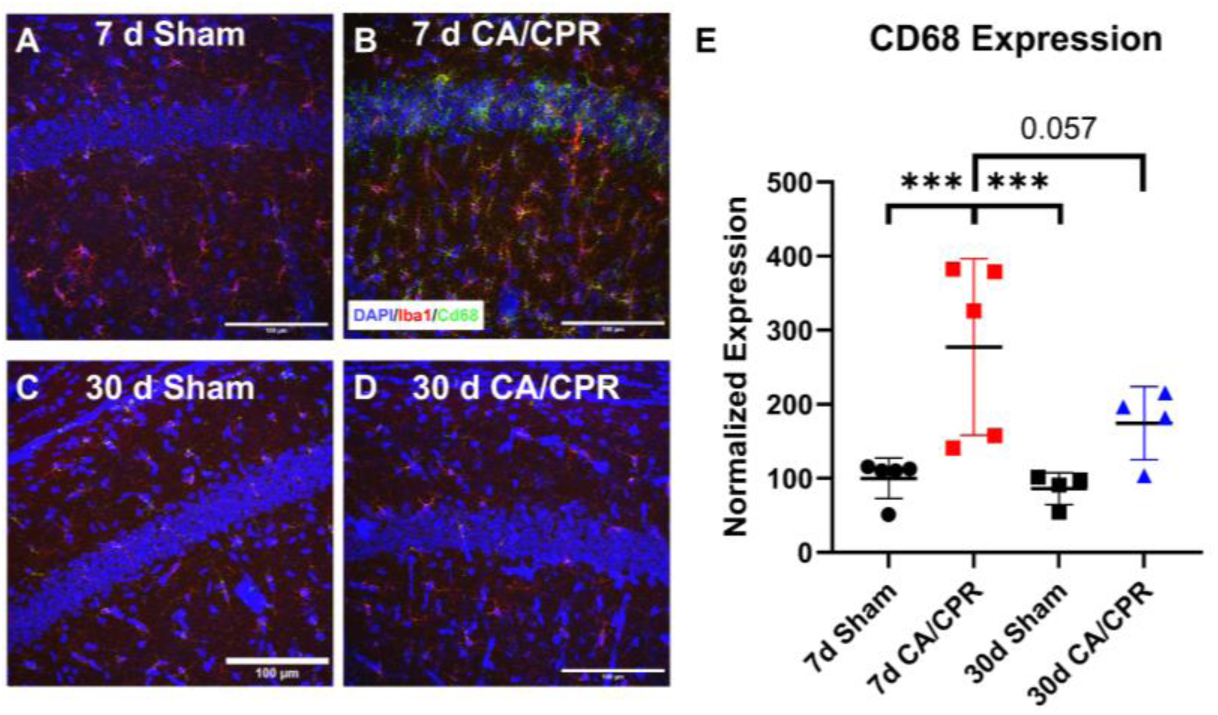
Protein expression levels of CD68 mirror the transcriptional data. Representative images of DAPI (blue), Iba1 (red), and CD68 (green) staining in the CA1 region of the hippocampus in 7d Sham animals (A), 7d CA/CPR animals (B), 30d Sham animals (C), and 30d CA/CPR animals. Quantification of protein expression of CD68 from confocal images (E). (*** = p < 0.001, GLMM).

### Protein expression levels of CD68 correspond with the GCI induced changes observed in mRNA transcripts

Changes in transcript levels do not always correlate with changes in the protein levels, thus it was necessary for us to confirm the results we obtained from the transcriptional analysis. Quantitative immunohistochemistry for CD68 expression was performed either 7- or 30-days after sham or CA/CPR surgery. Consistent with the transcriptional analysis, we found that there is a significant increase in CD68 expression 7 days after CA/CPR (227.2 ± 119.1% of 7d sham; p < 0.05). There was also recovery to near 30-day sham levels (86.12 ± 21.58% of 7d sham) in 30 -day CA/CPR animals (174.5 ± 49.38% of 7d sham; p = 0.057). This did not reach statistical significance, but the expression level is so different from 7d CA/CPR CD68 levels that it is reasonable to say there is recovery of protein levels of CD 68 that is consistent with the transcriptional analysis.

### Endogenous Recovery of GCI- Induced Hippocampal LTP Dysfunction is Diminished by Post-CA/CPR Ovariectomy

The timeline of LTP recovery corresponds with the onset of sexual maturity in female mice (PND 48-53 at time of recording)^48^, an observation that suggests a link between sex hormone signaling and endogenous recovery of hippocampal LTP after GCI. We have previously shown that there is negligible circulating estradiol in female mice until PND 28, reaching adult levels by PND 52^30^. Thus, our experimental design induces GCI during a developmental timepoint when estradiol is undetectable (PND 25) and measures functional outcome when there is abundant circulating estradiol (PND 48-53). Therefore, we next sought to discover how the surge in gonadal sex hormones that occurs during the onset of puberty is influencing hippocampal function after GCI.

To evaluate our hypothesis of gonadal sex hormone mediated recovery of hippocampal LTP in juvenile female mice subjected to GCI, we performed the CA/CPR or sham surgery between PND 21-25, followed by OVX 5-7 days later. Consistent with previous reports ^49^, the ovariectomy procedure itself, in sham animals, had no effect on hippocampal LTP measured 30-days after GCI (159.2±31.87% of BL, n=8) (Figure 5A and B). However, in animals that underwent the CA/CPR surgery, ovariectomy prevented the endogenous recovery of hippocampal LTP that occurs at 30-days post-surgery (123.2±18.95% of BL, p < 0.05 vs sham OVX, n=6) (Figure 5A and B). OVX without GCI also resulted in a decrease of the PPR (1.502±0.19, n=7) (Figure 5C) when compared to the value of the PPR in sham animals (1.734±0.2, n=7) (Figure 1C). OVX following CA/CPR resulted in a further reduction of the PPR (1.205±0.07, p < 0.01 compared to OVX alone, n=5) (Figure 5C). These changes in PPR suggest that gonadal hormone signaling modifies pre-synaptic release probability in juvenile female mice and that circulating sex-hormones are involved in the recovery of hippocampal LTP after CA/CPR. However, GCI or gonadal sex hormone induced changes in PPR do not correlate with impairments and endogenous recovery of hippocampal LTP.

**Figure 5:**
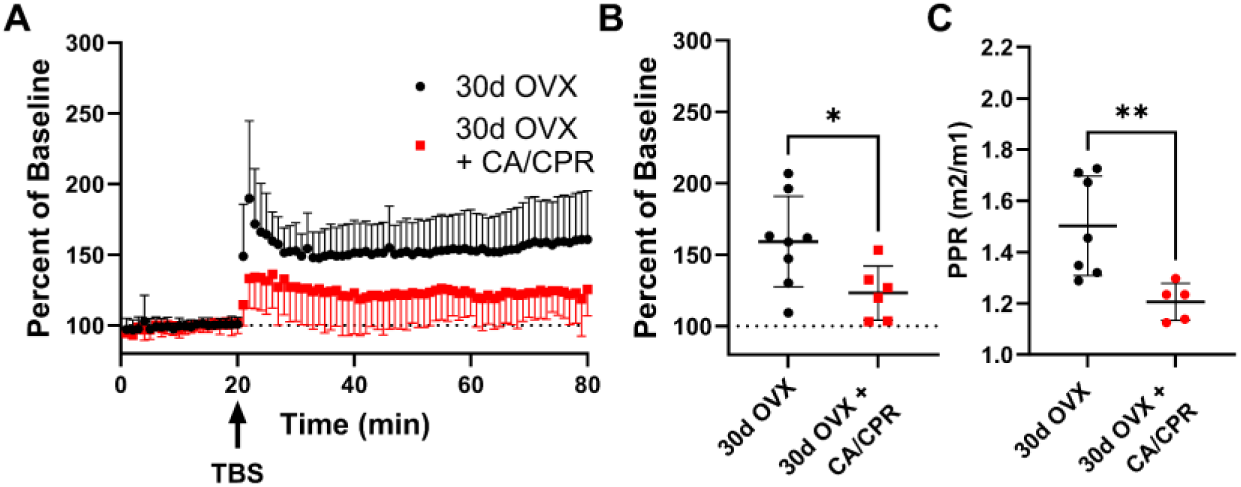
Ovariectomy after CA/CPR prevents endogenous recovery of LTP. (A) Animals that were ovariectomized after CA/CPR remain dysfunctional in the ability to undergo hippocampal LTP, relative to ovariectomized control animals, 30-days after CA/CPR. (B) Quantification of the last 10-minutes of the recordings performed in (A). (* = p < 0.05, Students T test). (C) Paired Pulse Ratio is not influenced by ovariectomy alone but is reduced by CA/CPR. (** = p < 0.01, Students T-test).

### Chronic Microglia Mediated Neuroinflammation Underlies Hippocampal LTP Deficits in Ovariectomized Female Mice Subjected to CA/CPR Induced GCI

We have shown that the developmental onset of gonadal sex hormone signaling in juvenile female mice profoundly effects the functional outcome of the hippocampus in survivors of early life GCI. However, the mechanism by which gonadal sex hormone signaling is mediating the endogenous recovery of hippocampal plasticity is less clear. To further understand the mechanism of endogenous hippocampal plasticity recovery after GCI and the lack of recovery from GCI after OVX we included a group of mice that underwent GCI and subsequent OVX in our transcriptional analysis. Upon further examination of this transcriptional data, we found that Cd68 expression remains significantly elevated 30-days after CA/CPR in OVX animals (n = 5) relative to CA/CPR animals with intact circulating hormones (n = 4) and sham animals (n = 3) (Figure 6A). This pattern of CD68 expression corresponds with the observed pattern of LTP dysfunction after GCI (Figure 1 A): the dysfunctional LTP observed 7-days after GCI corresponds with increased expression of CD68 and animals that undergo GCI with OVX also have increased expression of CD68 30-days after GCI, whereas sham animals or gonadally intact 30-day CA/CPR survivors do not have elevated expression of CD68. We also found that the mRNA expression of a pre-synaptic calcium channel, Cacna1b (N-type calcium channel), corresponds with the pre-synaptic plasticity dysfunction we observed after GCI or OVX (Figure 1 C and Figure 5C). These results further implicate neuroinflammatory processes in LTP impairments and hormone-dependent recovery after GCI and also highlight the mechanistic differences of GCI induced dysfunction of pre-and post-synaptic compartments. Post-synaptic plasticity is altered by GCI and modulated by gonadal sex hormones, whereas pre-synaptic plasticity is also modulated by GCI and gonadal sex hormones but does not endogenously recover after GCI and is not influencing LTP after GCI.

**Figure 6:**
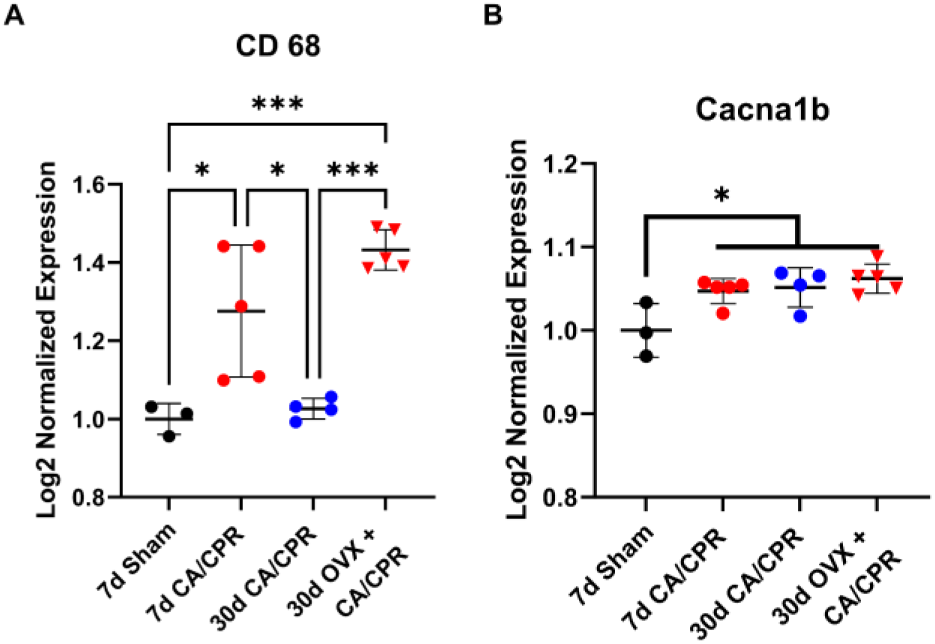
GCI induced transcriptional changes associated with GCI induced post-and pre-synaptic plasticity alterations. (A) CD68 mRNA is increased 7 days after CA/CPR with a recovery of expression to sham levels by 30 days post CA/CPR. Removal of gonadal sex hormones allows CD68 expression levels to remain elevated 30 days after CA/CPR. (* = p < 0.05, ** = p < 0.001, One-way ANOVA) (B) N-type calcium channel transcripts are significantly elevated by CA/CPR at all time points despite recovery of hippocampal LTP. (* = p < 0.05, One-way ANOVA).

### Signaling Through ERα Estrogen Receptors Mediates the Endogenous Recovery of Hippocampal LTP After GCI

Female gonadal sex hormones elicit their actions through various receptor signaling pathways ^50^. Therefore, we decided to test the possibility that activation of estrogen receptors with E2, or the receptor specific agonists would recover LTP in ovariectomized females. Animals underwent the CA/CPR procedure and were ovariectomized 5-days later with hormone replacement via sub-cutaneous osmotic pumps to deliver 1mg/kg/day of either vehicle, estradiol (E2), ERβ agonist (DPN), or ERα agonist (PPT). As expected, females that underwent CA/CPR and OVX+vehicle administration had diminished hippocampal LTP (127.9±21.23% of BL, n=8) 30-days later. Chronic administration of DPN or E2 in OVX animals yielded moderate, but not significant, recovery of hippocampal LTP (145.8±24.79% of BL for E2, n=11, 160.1±37.74% of BL for DPN, n=8) 30-days after CA/CPR. Chronic administration of PPT, however, did yield a significant recovery of LTP (170.4±41.77% of BL, n=12) (Figure 7A and B). To demonstrate the efficacy of ERα receptor activation to mediate an acceleration of recovery of hippocampal LTP after GCI we also evaluated the effect of acute PPT treatment on hippocampal LTP. Juvenile female mice were I.P injected with PPT (10mg/Kg) 6-days after CA/CPR and LTP recordings were performed 1-day later. When compared to 7-day CA/CPR vehicle injected animals (107.9 ± 18.85, n = 7), acute PPT treatment significantly recovered hippocampal plasticity (157.3±36.7%, n=6) (Figure 7 D and E). This effect on hippocampal LTP, by either chronic or acute PPT treatment, was not accompanied by recovery of the PPR (Chronic Treatment: 1.41±0.16, n=11; Acute Treatment: 1.45±0.13, n=6) (Figure 7 C and F), however E2 administration did increase the PPR (1.605±0.2, n=7) to near sham levels (1.734±0.2 Figures 1c and 5C, n=7) without a concomitant recovery of hippocampal LTP. This result suggests that ERα receptor signaling mediates endogenous recovery of hippocampal LTP in juvenile female mice after CA/CPR induced GCI without modifying pre-synaptic short-term plasticity.

**Figure 7:**
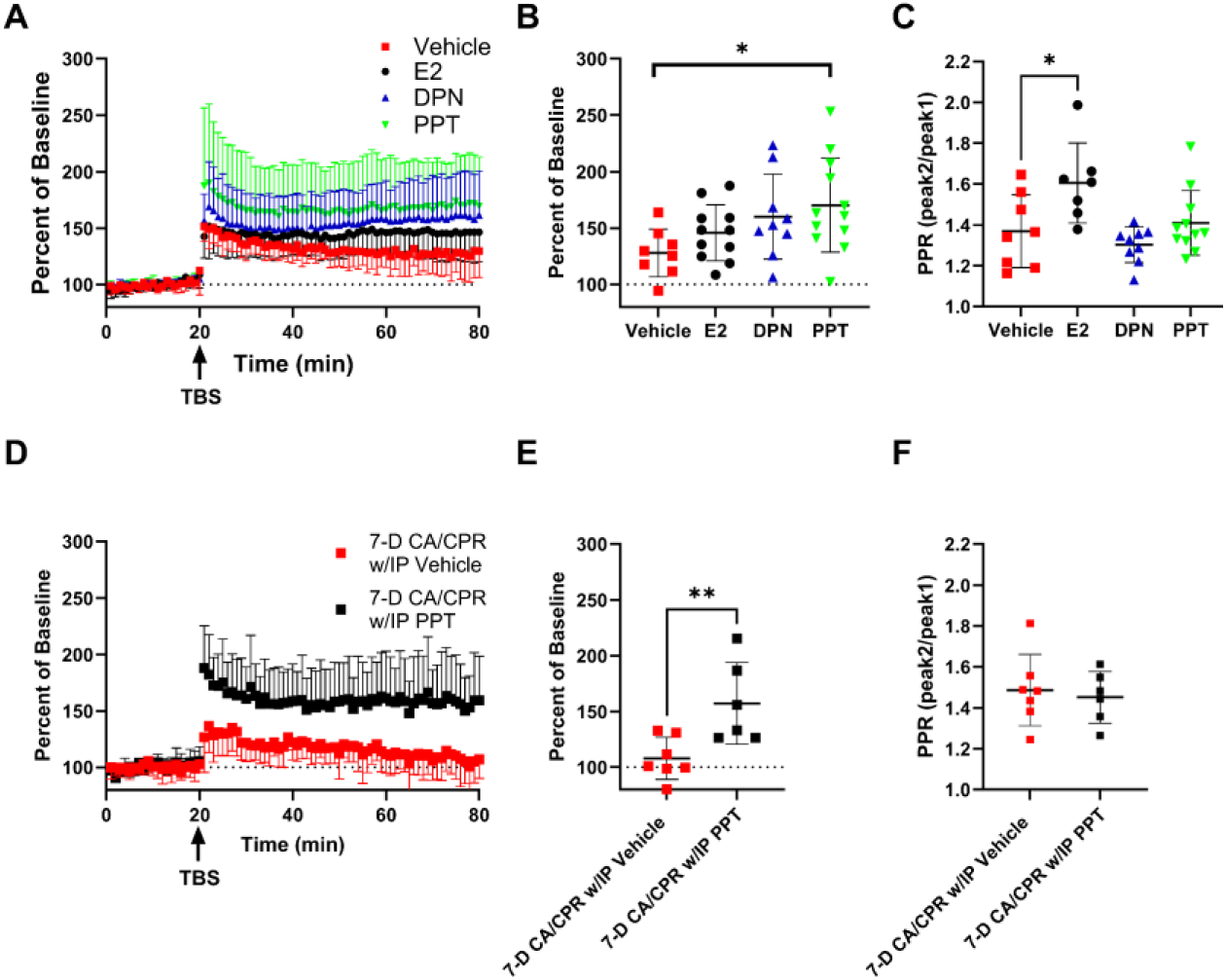
Endogenous Recovery of hippocampal LTP in juvenile female mice is mediated by ERα receptor signaling. (A) Hippocampal LTP recordings performed 30-days after CA/CPR in ovariectomized animals that were continuously treated with either vehicle, E2 (Estradiol), DPN (ERβ agonist), or PPT (ERα agonist), via subcutaneous osmotic pump. (B) Quantification of last 10-minutes of LTP recordings in (A). (* = p < 0.05, One-way ANOVA). (C) Paired pulse ratio measurements from recordings performed in (A) (* = p < 0.05, One-way ANOVA). (D) Hippocampal LTP recordings performed 7-days after CA/CPR. Black squares represent recordings from animals 7-days after CA/CPR and treated with 10mg/Kg PPT 6-days after CA/CPR. (E) Quantification of last 10-minutes of LTP recordings in (D). (** = p < 0.01, Student’s T-Test). (F) Paired pulse ratio measurements from recordings performed in (D).

## Discussion

Our study is the first to utilize a translational mouse model of juvenile global cerebral ischemia to assess hippocampal function at chronic timepoints after the ischemic insult in juvenile female mice. We have found that GCI in juvenile female mice results in a transient impairment of hippocampal function and endogenous recovery that occurs on a timeline that corresponds with the onset of sexual maturity (PND 48-53 at time of recording), suggesting a link between sex hormone signaling and the observed recovery of hippocampal LTP and spatial learning and memory. Contrary to adult animals ^17^, these results demonstrate the unique ability of the juvenile female hippocampus to endogenously recover function after an early life ischemic injury. LTP recovery was attenuated in females subjected to ovariectomy after acute cell death processes have subsided (5-7 days post-injury), suggesting the surge of gonadal sex hormones that occurs at the onset of sexual maturity in juvenile female mice mediates endogenous recovery of hippocampal LTP after CA/CPR, independent of alterations in neuronal cell death. Using agonist-selective hormone replacement, we implicated ERα as the primary mediator of sex-hormone dependent recovery of LTP. This recovery of hippocampal function highlights the need to further study the mechanism of recovery of hippocampal function in juvenile animals of both sexes. This mechanistic insight may help identify therapeutic targets which can enhance hippocampal function in adult survivors of CA/CPR and accelerate recovery of hippocampal function in juvenile CA/CPR survivors during the critical school-aged years.

Hippocampal LTP is widely accepted as the cellular mechanism that underlies declarative learning and memory abilities ^51, 52^. LTP in the hippocampus has been strongly associated with cognitive function and disruption of LTP greatly diminishes the ability to learn and remember ^53^. Our data support this, as we showed that in addition to the GCI-induced deficit and recovery of hippocampal LTP, there is also a deficit and recovery of spatial learning and memory after GCI. Therefore, we used this measure of hippocampal function as a surrogate for learning and memory ability in juvenile female mice that survive CA/CPR. The hippocampus is a brain region that is particularly sensitive to ischemic insult and our lab has previously shown that there are significant and similar levels of neurodegeneration in the hippocampus of adults and juveniles, regardless of sex ^7, 11, 18, 19, 54^. Thus, despite similar injury, juveniles have a remarkable ability to regain function in the surviving circuitry of the hippocampus. While it is possible that neuronal replacement could contribute to this recovery of function, we have previously reported that neurogenesis is not increased in juvenile male mice after GCI ^19^. Interestingly, BDNF levels correlate with LTP and associated learning and memory impairments in juvenile male mice subjected to CA/CPR, with levels decreasing at 7 days and recovering to sham levels at 30 days. However, contrary to males, our transcriptional analysis revealed a non-significant reduction in BDNF levels at 7 days in juvenile females after CA/CPR (Figure 1 D). This data suggests that while impairment and endogenous recovery occur in both sexes, there are distinct mechanisms underlying these changes juvenile females and males.

Because the mechanisms of endogenous recovery of hippocampal LTP after GCI remained largely unknown, we incorporated transcriptional analysis using NanoString technology to provide additional insight. We observed many GCI induced transcriptional changes but focused on those whose expression correlated with the LTP deficit at 7-days and recovery at 30-days. Therefore, we incorporated an analysis pipeline that excluded transcripts that are developmentally regulated and allowed us to identify those that recover by 30-days post GCI. Of these, Cd68 was the most highly GCI-regulated transcript in our assay that met these criteria. Cd68 is highly upregulated in activated microglia and is associated with microglial phagocytosis of synapses and cellular debris ^55, 56^. Interestingly, Cd68 expression in activated microglia is also modulated by estrogen signaling in female rodents ^57^. Also, several recent studies have implicated microglia mediated neuroinflammatory processes in the disruption of LTP in the SC-CA1 circuit^58–60^ and Varghese et al., 2023 have shown that LTP deficits, which result from brain injury, can be attenuated by partial depletion of microglia^61^. Thus, we confirmed the transcriptional result by performing CD68 IHC. Taken together, our results suggest a scenario where GCI in pre-pubescent females induces an increase in neuroinflammatory processes which are disruptive to hippocampal LTP and learning. The surge of gonadal hormones that occur at the onset of puberty modulates this neuroinflammatory response, dampening it, and once again allowing for the induction of hippocampal LTP and spatial learning. Removal of endogenous sex hormone signaling during this time prevents endogenous recovery of hippocampal LTP and promotes a chronic neuroinflammatory state. However, CD68 is also expressed on circulating macrophages, suggesting further work is needed to clarify cellular source of neuroinflammatory response impairing LTP in our model. It will also be highly informative for future studies to assess neuroinflammation in adult female GCI populations, where endogenous recovery of hippocampal function does not occur.

Extensive evidence exists which suggests that female sex hormones, in particular estrogen, are neuroprotective in many models of brain injury and memory related pathologies ^24, 25^. However, the study of sex hormone signaling is complex and multifaceted. E2 is reported to have fast non-genomic effects on hippocampal function as well as more prolonged genomic effects ^62^. Expression patterns of estrogen receptors (ERα, ERβ, and GPR 30) are known to differ between the sexes and are also differentially expressed between brain regions and cellular compartments (membrane vs cytosolic vs nuclear localization)^63–65^. The benefits of estrogen replacement therapy on cognitive function are also variable, depending on dose, duration of treatment, and the time that has elapsed since endogenous sex hormone signaling was disrupted (via hysterectomy or menopause) and start of therapy ^66–68^. In regard to learning, memory, and plasticity, there are reports of ERα and ERβ having antagonistic actions (one ER disrupting the actions of the other), being cooperative (one ER enhancing the activities of the other), or ratio metric (the ratio of expression levels of ERα and ERβ determine the functional outcome of their activity)^22, 29, 69^.

Thus, in addition to identifying the necessity of gonadal sex hormone signaling to endogenous recovery of hippocampal LTP after GCI, we also sought to further understand how individual estrogen receptors (ERα or ERβ) are influencing hippocampal LTP after GCI. We determined that OVX alone did not impair LTP in female mice, but OVX did prevent the endogenous recovery of hippocampal LTP following CA/CPR. OVX after CA/CPR also resulted in elevated CD68 transcript levels 30 days after CA/CPR in female mice, further supporting a role for neuroinflammation in LTP impairments after GCI. We also found that the ERα agonist PPT was able to significantly restore LTP in OVX females while DPN, the ERβ agonist, did not, implicating ERα signaling in the endogenous recovery of LTP. Surprisingly, E2 replacement itself did not significantly promote recovery of hippocampal LTP which could be due to the pre-pubescent developmental timepoint at which the E2 replacement therapy was started. We did however provide evidence that acute pre-pubescent activation of ERα receptors is able to recover hippocampal plasticity, demonstrating the potential of targeting ERα receptors to speed recovery of hippocampal LTP after pediatric GCI. Overall, we suggest that after GCI in juvenile females, endogenous recovery of hippocampal LTP is mediated by ERα signaling. Importantly, our experimental design differentiated the neuroprotective effects of estrogen from their neurorestorative effects, indicating a new restorative function for the hormonal surge observed in adolescence. Additional studies will be needed to determine if this acute effect of ERα activation on hippocampal function is long lived or is applicable in juvenile males. It will also be highly informative for future studies to perform transcriptional analyses in adult females after GCI to assess whether chronic neuroinflammation is underlying the lack of recovery of hippocampal function in this population. Future studies should also examine the effect that ERα signaling has on neuroinflammation in both adult and juvenile female animals.

Accompanying the observed hippocampal LTP deficit, we also observed a GCI-induced chronic increase in the probability of neurotransmitter release (reduced PPR) after GCI (Figure 1 C, Figure 5 C, Figure 7 C and F) which was not reported in juvenile male animals ^19^. Our data demonstrates that the mechanisms of pre-synaptic and post-synaptic dysfunction after GCI are likely unrelated and that presynaptic functional changes after GCI may be sexually dimorphic. The probability of neurotransmitter release is dependent on pre-synaptic calcium signaling and modulated by local calcium concentrations in the pre-synaptic bouton ^70^. Thus, we propose a mechanism of GCI-induced increase in pre-synaptic calcium signaling, resulting in increased probability of neurotransmitter release. This mechanism is supported by our transcriptional analysis, as we found a small, but significant, increase in the expression of Cacna1b (Ca_V_2.2) that is associated with the GCI induced reduction of the PPR (Figure 6B) and consistent with previous reports of this N-type calcium channels ability to modify pre-synaptic function ^71^. Our results also suggest that gonadal sex hormone signaling modifies pre-synaptic short-term plasticity in juvenile female mice. The absence of these gonadal sex hormones in ovariectomized females leads to increased neurotransmitter release at the SC-CA1 synapse, without disrupting LTP, further supporting the dissociation of pre-synaptic and post-synaptic mechanisms of plasticity^72^. However, the increased release probability we observed is not influencing SC-CA1 hippocampal LTP, as it remains elevated despite endogenous recovery of LTP. Pre-synaptic plasticity has been associated with learning and memory processes and our results highlight the need for future studies into how this phenomena may impact other aspects of circuit function beyond LTP ^73, 74^.

Collectively, we have described a mechanism of recovery of hippocampal function after CA/CPR induced GCI in juvenile female mice that is dependent on gonadal sex hormone signaling, in particular, activation of ERα. We provide evidence that GCI in juvenile females alters pre-and post-synaptic plasticity, but that there are differing underlying mechanisms of these alterations. We also provide evidence implicating neuroinflammatory processes as an underlying mechanism of the observed post-synaptic LTP and spatial learning deficits in juvenile females. We are beginning to unravel the mechanisms behind the juvenile brain’s unique ability to repair itself with the goal of harnessing these mechanisms in other populations of GCI survivors to promote neurorestoration of hippocampal plasticity.

## ACKNOWLEDGEMENTS

This work was supported by R01NS046072-S1 (J.V), 23PRE1020271 (J.V), F99NS135766 (J.V), R01NS046072 (N.Q. and P.S.H.), R01NS118786 (P.S.H), R01NS092645 (P.S.H.), KO8NS097586 (R.M.D). We’d like to acknowledge Dr. Traystman for his lasting contributions to the inception of this study.

## AUTHOR CONTRIBUTIONS

Conceptualization, J.V., N.Q., and P.S.H.; Formal Analysis and Investigation, J.V., J.E.O., V.N.C., R.M.D., Methodology, E.T., D.M., M.F., I.V., and N.C.; Writing – Original Draft N.Q., J.V.; Writing – Review & Editing, N.Q., P.S.H., and R.M.D.; Supervision, N.Q, and P.S.H.; Funding Acquisition J.V., N.Q., R.M.D. and P.S.H.

