## Supplementary Figure for "Endogenous recovery of hippocampal function following global cerebral ischemia in juvenile female mice is influenced by neuroinflammation and circulating sex hormones"

### Supplemental Materials

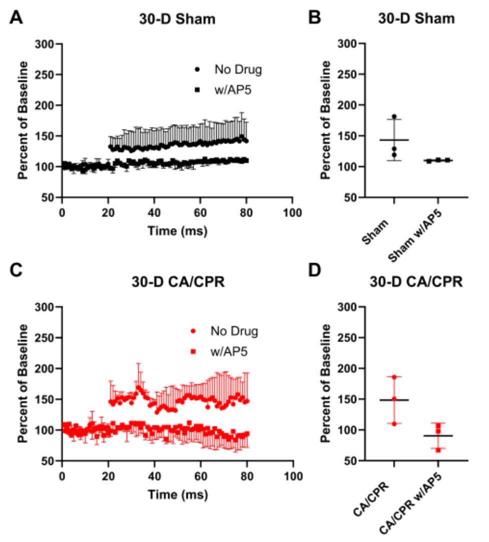


**Supplemental Figure 1: Hippocampal LTP relies on a post-synaptic mechanism.** Hippocampal LTP experiments performed in Sham (A) and CA/CPR (C) animals 30 days after surgery with or without AP5. Quantification of last 10 minutes of LTP in sham (B) and CA/CPR (D) animals.
